# Plant diversity in the diet of Costa Rican primates in contrasting habitats: a meta-analysis

**DOI:** 10.1101/2023.02.02.526906

**Authors:** Óscar M. Chaves, Vanessa Morales-Cerdas, Jazmín Calderón-Quirós, Inés Azofeifa-Rojas, Pablo Riba-Hernández, Daniela Solano-Rojas, Catalina Chaves-Cordero, Eduardo Chacón-Madrigal, Amanda D. Melin

## Abstract

In human-modified tropical landscapes, the survival of arboreal vertebrates, particularly primates, depends on their plant dietary diversity. Here, we assessed diversity of plants included in the diet of Costa Rican non-human primates, CR-NHP (i.e. *Alouatta palliata palliata*, *Ateles geoffroyi*, *Cebus imitator*, and *Saimiri oerstedii*) inhabiting different habitat types across the country. Specifically, we assessed by analyzing 37 published and unpublished datasets: (i) richness and dietary α-plant diversity, (ii) the β-diversity of dietary plant species and the relative importance of plant species turnover and nestedness contributing to these patterns, and (iii) the main ecological drivers of the observed patterns in dietary plant . Diet data were available for 34 *Alouatta*, 16 *Cebus*, 8 *Ateles*, and 5 *Saimiri* groups. Overall dietary plant species richness was higher in *Alouatta* (476 spp.), followed by *Ateles* (329 spp.), *Cebus* (236 spp.), and *Saimiri* (183 spp.). However, rarefaction curves showed that α-diversity of plant species was higher in *Ateles* than in the other three primate species. The γ-diversity of plants was 868 species (95% C.I.=829-907 species). The three most frequently reported food species for all CR-NHP were *Spondias mombin*, *Bursera simaruba*, and *Samanea saman*. In general, plant species turnover, rather than nestedness, explained the dissimilarity in plant diet diversity (β_sim_> 0.60) of CR_NHP. Finally, primate species, habitat type (life zone and disturbance level) and, to a lesser degree, sampling effort were the best predictors of the dietary plant assemblages. Our findings suggest that CR-NHP diets were diverse, even in severely-disturbed habitats.

## 1. Introduction

Dietary diversity and behavioral flexibility are key factors on the survival probability of most vertebrates (including humans) worldwide, because food availability frequently fluctuates spatio-temporally [1–3]. Food fluctuation is in human-modified landscapes is particularly notable, because non-sustainable anthropogenic activities (e.g. forest deforestation, hunting, urbanization) often transform large, diverse and heterogeneous forests into low-diversity and homogeneous poor-quality forest habitats [4, 5]. Habitat degradation can promote diverse feeding behavioral adjustments in vertebrates, including, diet diversification, diet supplementation with exotic and/or cultivated food species, changes in the foraging patterns and food preferences, temporal changes in time budget, inventions of new traditions (including the consumption of new food items), and others [6–9].

Among vertebrates, non-human primates (NHP) are one of the most eclectic or flexible taxawith respect to food use [2,10–12], a behavior that may allows them to adjust their diets under critical conditions such as human-modified and/or strongly seasonal forests (2,12-14).However, there is not comparative information about how primates with different dietary preferences (e.g. frugivores vs. folivores vs. omnivores) cope with seasonally or human-modified environments, in particular, how they diversify their plant food diet under such scenarios and which mechanisms drive this changes. Of particular concern are neotropical primates, given the high rate of forest conversion in their habitats and their high reliance on resources provided by forest tree species [14, 18]. Plant diet diversity can be divided into nested levels of organization to elucidate the mechanisms regulating the inclusion of plant species in primate diets, these division include: (i) α-diversity, or the diversity of dietary plant species included by a primate species in a given site, (ii) β-diversity, or the variation of dietary plant species composition among sites or dietary plant species dissimilarity, and (iii) γ-diversity or the total diversity of dietary plant species in a landscape or geographic area [23, 24]. Furthermore, dietary plant species β-diversity can be further studied as the result of two ecological mechanisms to explain the dissimilarity between plant species assemblages in primate diets: spatial species turnover and nestedness of species assemblages [25]. Spatial species turnover measures the replacement of plant species in one site by distinct species in the other site, and nestedness of species assemblages refers to the loss (or gain) of species in only one of the sites and, as consequence, the poorest site contains only a subset of the species composition of the richest one [25, 26]. The diversity of plants used as food sources by neotropical primates depends, among other factors, on aspects of their natural history, such as trophic guild, social organization, group size, local plant diversity and plant species-specific abundance. For instance, highly frugivorous primates living in fusion-fission communities such as spider monkeys (*Ateles* spp.) invest more time traveling and foraging on a more diverse array of plants compared with more folivorous primates, living in cohesive small groups, such as *Alouatta* spp. [27–30]. It has been hypothesized that, in contrast with mature leaves, the distribution of fleshy fruits (and immature leaves) is sparse in space and time, forcing animals to diversify their diets [17,27,30] and, in the particular case of *Ateles* spp., the social dynamics can increase the probability of finding new food resources, because they can forage in multiple subgroups in different regions, over a large area throughout the forest [28,31,32].

Costa Rica supports a large diversity of primates, with a total of four species and ca. six subspecies: mantled howler monkeys (*Alouatta palliata palliata*), spider monkeys (*Ateles geoffroyi frontatus*, *A. g. geoffroyi*, *A. g. ornatus*), squirrel monkeys (*Saimiri oerstedii citrinellus*, *S. o. oerstedii*), and capuchin monkeys (*Cebus imitator*). *Alouatta* is the most widely distributed primate in the country, inhabits a large variety of natural and human-modified habitats [33, 34], and lives in groups from <6 to ca. 45 individuals. As with most species in the genus *Alouatta*, its diet is often classified as folivorous-frugivorous [12,13,27]. *A. geoffroyi* is mainly restricted to continuous/large well-preserved forests, their troop sizes are >25 individuals, and its diet is strongly frugivorous [17,31,33]. *C. imitator* is distributed in a variety of continuous and fragmented habitats throughout the country. Capuchin groups vary from 8 to 35 individuals and its diet is classified as insectivorous-frugivorous [28,33,35]. Finally, *S. oerstedii* is an endemic species restricted to the central and southern Pacific coast of Costa Rica and part of the Chiriqui province, Panama. Their groups can range from 20 to >90 individuals and, as *C. imitator*, its diet is described as insectivorous-frugivorous [33,34,36].

Due to ongoing destruction of their natural habitats and the growing contact between these animals and humans, particularly in highly touristic near protected areas and/or urbanized areas throughout the country, e.g. [37], the former three species are considered at risk of extinction locally according to the Costa Rican environmental authorities [38], endangered or vulnerable across their former distribution [39]. However, to date, our knowledge on the diversity and flexibility of plant diet in Costa Rican non-human primates (hereafter CR-NHP) inhabiting contrasting habitats is poorly understood. There are a number of published and unpublished studies on the feeding behavior of CR-NHP, but no study has synthetized the available information on this topic to further understand the mechanism that drive diet species selection by these primates. This issue preludes any generalization countrywide about resource use by these locally threatened primates. Also, from the conservation point of view, constrains the design and implementation of appropriate management strategies for their conservation and their habitats. For instance, detailed information on the main plant food species used by CR-NHP might be crucial to develop efficient reforestation programs to improve connectivity between adjacent fragments inhabited by different primate populations, as has been proposed for *A. palliata* [40, 41].

In this study, we investigated, for the first time, the diversity of plant species in the diet of the four CR-NHP. For this, we reviewed countrywide published and unpublished information on feeding ecology of these animals. Our main objective was to describe and compare the diversity of plant species included in their diet under different habitats (i.e. diet flexibility). In addition, we explored several potential ecological drivers to explain the patterns found in this review. Specifically, we assessed: (i) the α and γ-diversity of plants in their diet and the most important plant food species for each primate species, (ii) the dissimilarity of food plant assemblage in the diet between the four CR-NHP (i.e. β-diversity), and the relative importance of food plant species replacement (i.e. species turnover) and food plant species loss (i.e. species nestedness) in the diet to explain these patterns, and (iii) the role of different habitats traits and CR-NHP species as predictors of the observed plant assemblages in the diet. Finally, we discuss the implications of the observed patterns on the conservation and management of the four CR-NHP and their habitats. We predict that the α-diversity of plants species in the diet of the most frugivorous primates will be higher (e.g. *A. geoffroyi)* than in the diet of less frugivorous and/or omnivorous primates (e.g. *A. palliata, C. imitator*, and *S. oerstedii).* Furthermore, because CR-NHP inhabit forest areas with contrasting plant compositions (e.g. Tropical Dry Forest and Tropical Rain Forest [42, 43]) and disturbance levels, we also expect that species turnover, rather than nestedness, will be the main ecological mechanism influencing the β-diversity patterns in plant diet.

## 2. Materials and Methods

### 2.1. Study Region

Although a comprehensive survey of habitats occupied by CR-NHP has not been assessed to date, we estimate that 25-50% of the forested area in Costa Rica (i.e. ca. 3.8 Mha: [44]) is inhabited by at least one CR-NHP (commonly *A. p. palliata:* Ó.M.C. pers. obs.). The studies we review for this meta-analysis were performed for a discrete number of private and public forest habitats distributed throughout the seven provinces of Costa Rica (Figure 1). Historically, most multi-year and long-term primate studies in Costa Rica have been focused in three regions: northern Costa Rica (e.g. sector Santa Rosa of the Área de Conservación Guanacaste, Hacienda La Pacífica, and Palo Verde National Park, Guanacaste), the central Pacific coast (e.g. Manuel Antonio National Park, Puntarenas), and southern Pacific coast (e.g. Península de Osa, Puntarenas; Figure 1). Overall, these areas cover two main Holdridge’s Life Zones HLZ [43, 45]: Tropical Dry Forest (TDF) and Tropical Rainforest (TRF). In addition, some studies were conducted in three other HLZs: Tropical Lowland Rainforest (TLRF), Humid Premontane Forests (HPF), and Tropical Wet Forest (TWF). Throughout the study region, primates inhabit a large range of private forest fragments (ranging from <10 ha to 250 ha) under different successional stages and areas of continuous forest located within national parks (Figure 1).

**Figure 1.**
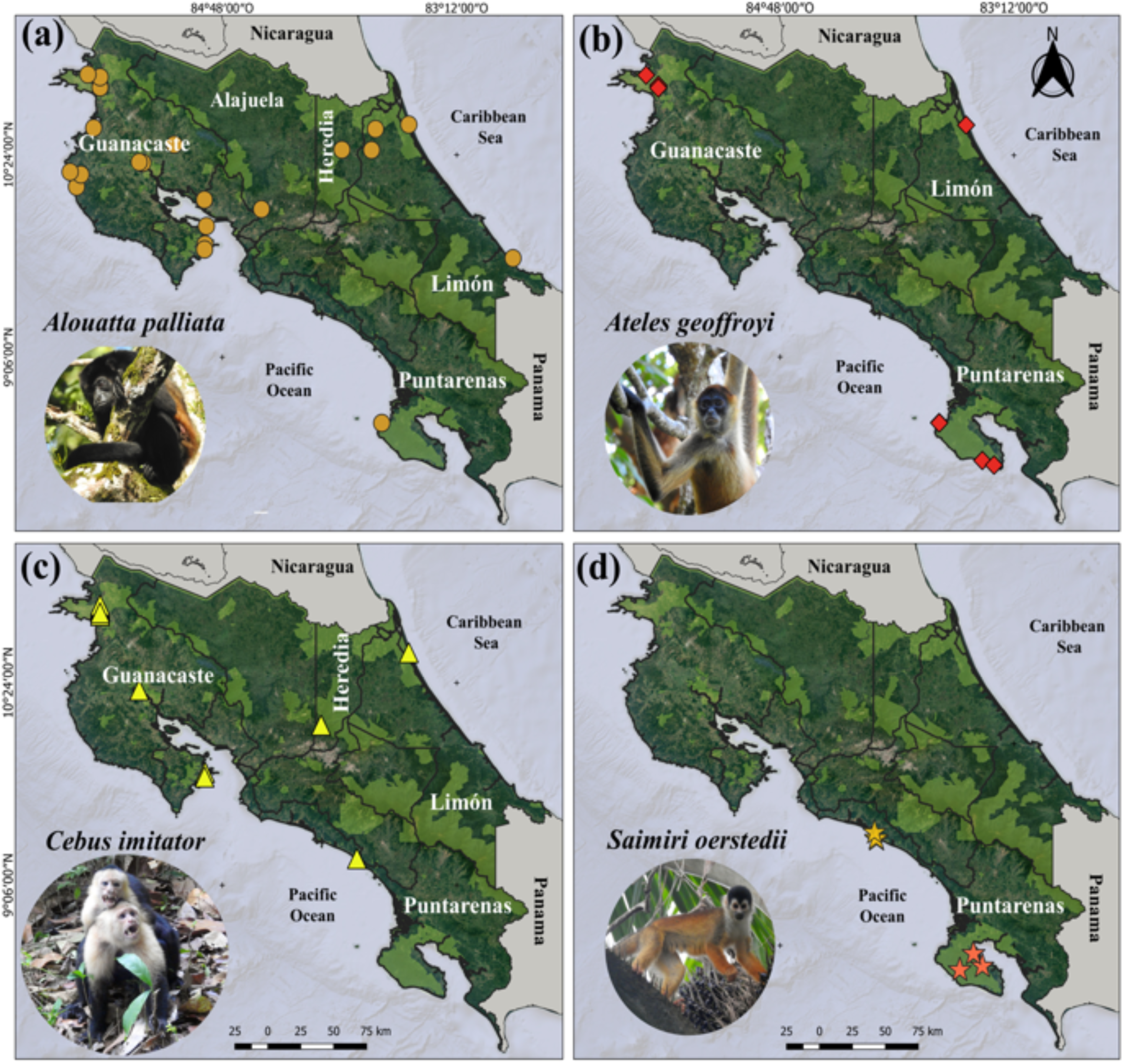
Distribution of the study sites for each primate species reported by the studies included in the review. The exact location of forest fragments for *Alouatta palliata palliata* (a), *Ateles geoffroyi* (b), *Cebus imitator* (c) and *Saimiri oerstedii* (d) are indicated with different symbols and colors. Pale green polygons represent the protected areas of Costa Rica. Photos by Ó.M. Chaves. The first figure layer is a free Google Earth® satellite image.

The tropical dry forests of Guanacaste represent a large extension of coastal lowlands with a marked climatic seasonality [45, 46] and, typically, they are comprised of a complex mosaic of pastures, agricultural lands, forest remnants of different sizes and successional stages, and scattered human settlements or small touristic cities [46] where the negative interactions with humans and CR-NHP are frequent (e.g. electrocution of howler monkeys in power lines: [41, 47]). The dry season starts in late November and extends throughout late April, while the rainy season extends from early May to November. Most of the canopy trees are deciduous and in the dry season, many species are defoliated. The annual average precipitation is ca. 1975 mm (range= 900-2400) and the average annual temperature is ca. 26 °C (range= 23-36 °C). Conversely, seasonality is low or moderate in the other four HLZs. Annual average precipitation can range from >3300 mm to >5500, and the average annual temperature is ca. 28 °C (range= 24-34 °C). Because of the scarcity of studies in TLRF, TWF, and HPF (see Table 1) and the similarities between these HLZ and the TRF in terms of climatic conditions (particularly the absence of a marked seasonality), in the analyses of plant *β-*diversity and the PERMANOVA described below, we pooled these four HLZ in a single category named “Rainy Forests”.

**Table 1.**
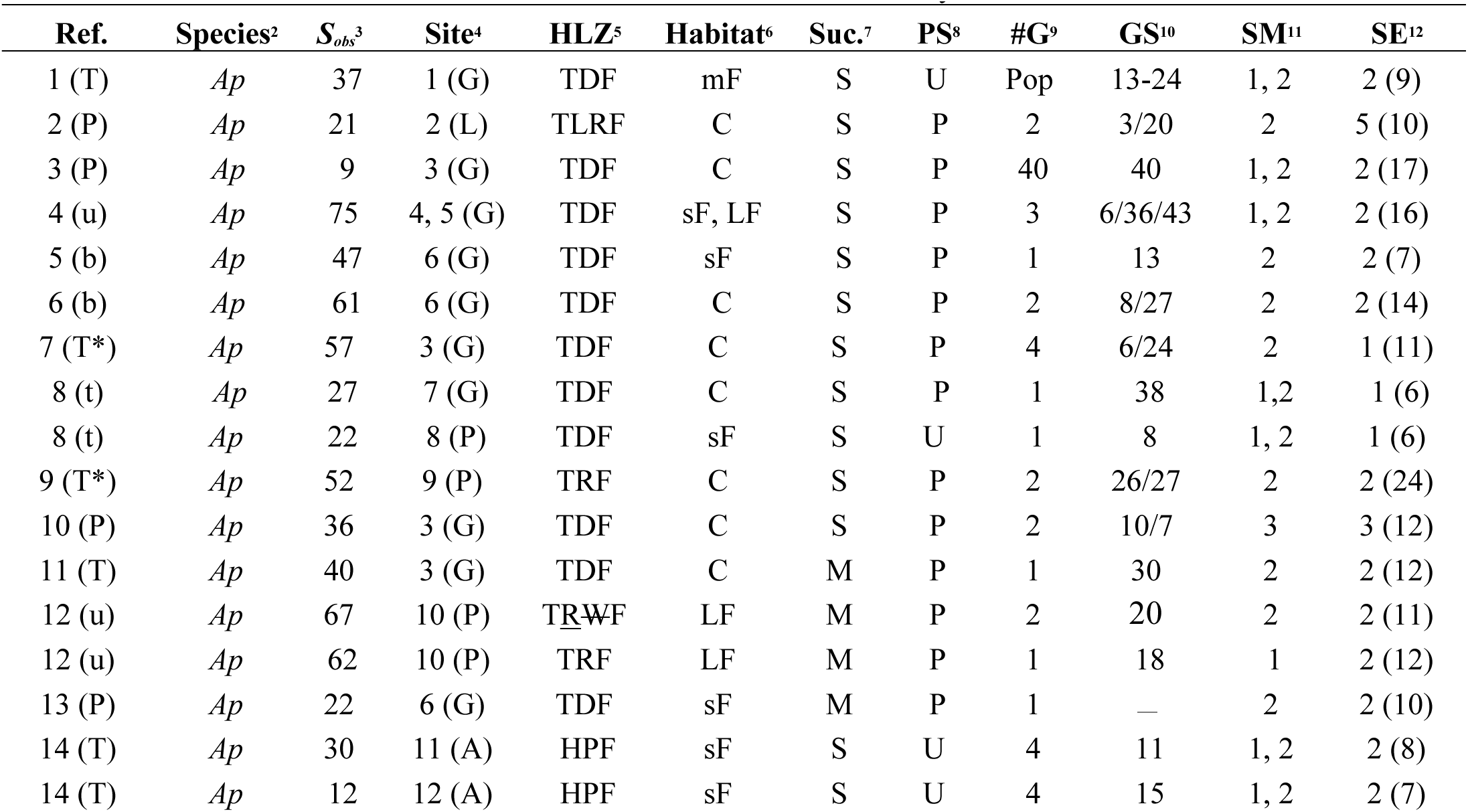
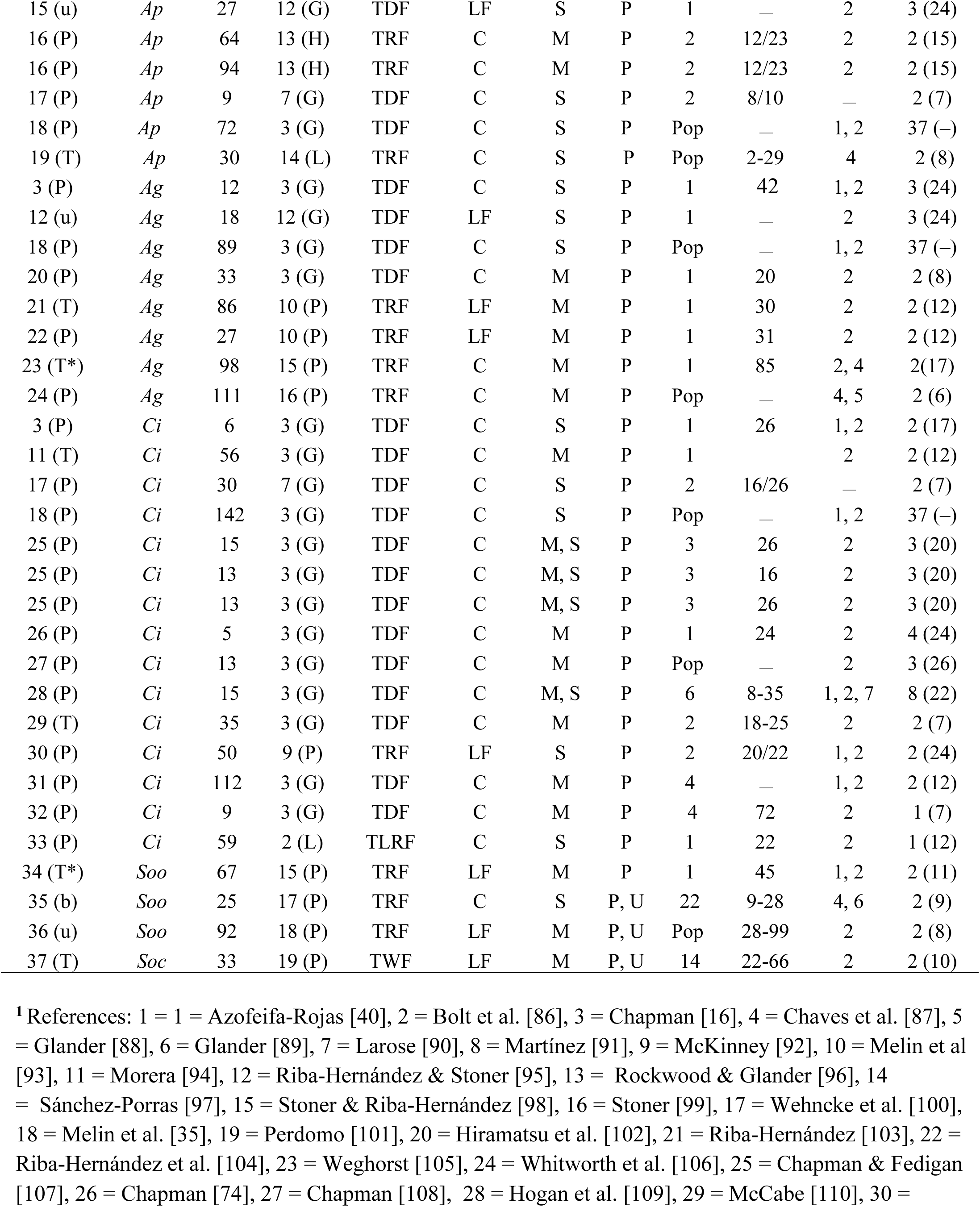
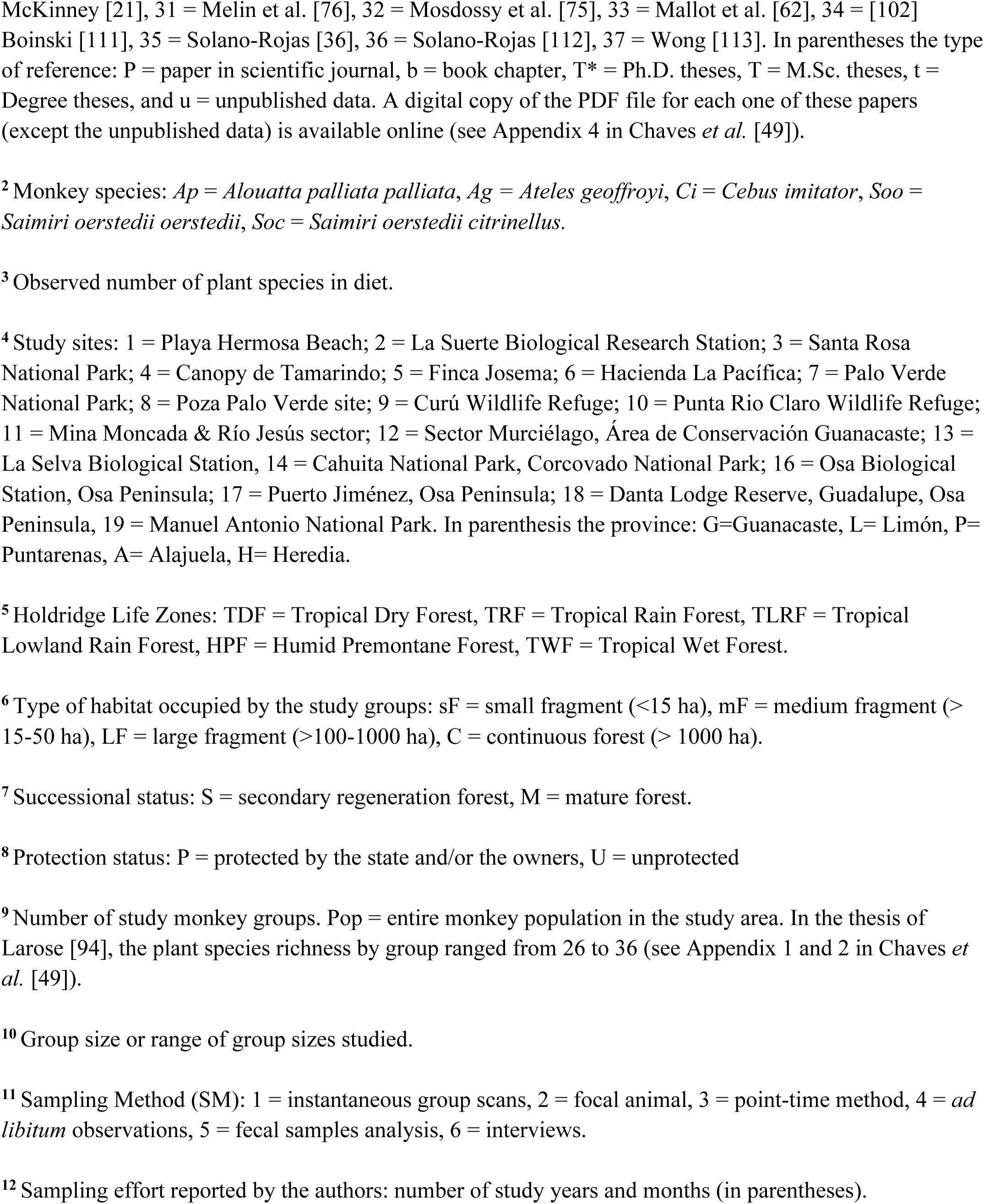
Studies on the diet of CR-NHP considered in the meta-analyses.

### 2.2. Literature Review and Data Collection

During a 9-mo period (from January to September 2022) we searched for all published and unpublished information using two general online databases: Google Scholar (https://scholar.google.com), and Web of Science (https://mjl.clarivate.com). Furthermore, to access dissertations from Costa Rican universities, we used the Kerwa repository from the Universidad de Costa Rica (https://www.kerwa.ucr.ac.cr/) and Biodoc repository from the Universidad Nacional (https://repositorio.una.ac.cr/). We restricted literature search to the period of 1975-2022 because references before 1975 are very scarce and rarely available online. We used specific keywords such as ‘Costa Rican primate diet’, ‘diet of Costa Rican primates’, ‘feeding behavior of Costa Rican primates’, and several combinations of keywords including common and scientific names of primates such as ‘diet + *Saimiri oerstedii* + squirrel monkeys + Costa Rica’. We also used the same keywords in Spanish to improve the probabilities to find dissertations and/or papers from local researchers.

Due to the relative scarcity of literature on the topic, we included diverse information sources such as: (i) papers published in peer-reviewed scientific journals, (ii) book chapters from recognized editorials, (iii) academic dissertations (including undergraduate and graduate theses), and (iv) technical reports. We also included unpublished data provided by coauthors of this meta-analysis. We only included field studies on wild, free-ranging monkey groups using standardized observational methods (e.g. instantaneous scans, focal animal, and *ad libitum* observations:[48]) and with a sampling effort ≥ 6 study months in our statistical analyses. However, to provide a comprehensive resource, datasets that did not fulfill these discrimination criteria were included in the online meta-analysis dataset (see Appendix 1 and 2 in Chaves *et al.* [49]).

Overall, we found 43 published and unpublished datasets on the diet of one or more CR-NHP (see Appendix 1 in Chaves *et al.* [49]). Most of these were scientific papers (58%), followed by M.Sc. theses (16.3%), Ph.D. theses and unpublished data (9.3% each). Commonly, these studies were focused on one or two different primate groups (range= 1-8 groups). In total, data on diet was available for 34 *Alouatta*, 16 *Cebus*, 8 *Ateles*, and 5 *Saimiri* groups (see Appendix 2 in Chaves *et al.* [49]). However, only 37 datasets met our criteria to be included in the statistical analyses (Table 1, see also Appendix 3 in Chaves *et al*. [49]).

### 2.3. Description of Field Data Collection Methods in the Literature Reviewed

Feeding behavior of howler monkeys *Alouatta palliata palliata* (hereafter *Ap*) was frequently studied in the TDF of Guanacaste and TRF of Puntarenas provinces, throughout the Pacific coast of Costa Rica. Overall, 1-8 free-ranging *Ap* groups previously habituated to human presence were observed in each study. However, most authors did not indicate the exact diurnal period they monitored, or the number of hours sampled per day (see Appendix 1 in Chaves *et al.* [49]). The most frequently used sampling methods for *Ap* were the focal animal method, with behavioral samples of 10 min, and instantaneous scan sampling at 15 min intervals [48]. However, some authors also used *ad libitum* observations [48], or a combination of two different methods (Table 1, see also the Supplementary Material). The study durations varied between studies, ranging from 6 to 33 months (Table 1).

The diet of spider monkeys *Ateles geoffroyi* (hereafter *Ag*) was collected from five different HLZ (i.e. TDF, TRF, TLRF, TWF, and PHF) in 3 out of 7 provinces of the country (Table 1). In 83% of cases, authors used the focal animal method with focal samples ranging from 2 to 10 min, but instantaneous scan sampling or a combination of *ad libitum* method plus collection and analyses of fecal samples were also used. Typically, the study durations ranged from 6 to 36 months, with an outlier that compiled data from 28 years in Santa Rosa National Park [35] (Table 1).

In contrast to studied on *Ap* and *Ag*, 82% of studies on the diet of the capuchin monkeys *Cebus imitator* (hereafter *Ci*) were performed in TDF (particularly Santa Rosa National Park), with the rest in TRF and TLF. The authors used the focal animal method or a combination of this method with group scans and the sampling effort typically ranged from 7 to 33 months, with one study reporting data across 37 years [35], (Table 1).

Squirrel monkeys *Saimiri oerstedii* are not found in the TDF in Costa Rica. The diet of these monkeys was studied in two regions in the central and southern Pacific coast of the country using a combination of field methods: *S. citrinellus* (hereafter *Soc*) was studied in a TWF (i.e. Manuel Antonio National Park) using focal and group scan methods, while *S. o. oerstedii* (hereafter *Soo*) was studied in Peninsula de Osa’ TRF using the group scan method and *ad libitum* observations. For both squirrel monkey subspecies the sampling effort was ≤11 months (Table 1).

Because the vast majority of studies included in this meta-analysis did not provide details on the number of feeding records or time devoted by each monkey group to the different food plant species (or if provided, the data were incomplete, i.e. 10 studies, see Appendix 2 in Chaves *et al.* [49]), it was not possible to determine the ‘top plant food species’ in the diet of the CR-NHP (i.e. those species that together represent ≥ 80% of feeding records: [32]). Importantly, no study provided this information for *So.* Instead, we determined the ‘plant species most frequently reported in the diet’ for each CR-NHP. For this, we organized the list of food plant species by the total number of monkey groups/populations (considering all the study sites for each CR-NHP) that exploited each plant species *i* (see Appendix 4 in Chaves *et al.* [49]). Finally, because the reviewed studies came from many researchers, most of whom were not specialists in plant taxonomy, and cover a wide time period (i.e. from 1972 to 2022), we detected diverse taxonomic inconsistencies, grammatical errors, and outdated classification with the reported plant scientific names (see Appendix 2 in Chaves *et al.* [49]). To prevent potential problems of over and/or underestimation of the plant species richness reported in each study, we standardized scientific names of the plants using the taxonomic name resolution service provided by Botanical Information and Ecology Network (https://bien.nceas.ucsb.edu/bien/tools/tnrs/), and the plant scientific names were updated using Tropicos plant database (https://www.tropicos.org/).

### 2.4. Statistical Analysis

#### 2.4.1. Plant α-Diversity of Diet

We analyzed patterns of dietary plant diversity (i.e., including both species and morphospecies) of CR-NHP using Hill numbers (*q*), metrics that represent true diversities because they obey the replication principle [50]. Hill numbers are expressed in ‘units of species’ which can be plotted in a single graph to compare the diversity profiles as a continuous function of the parameter *q* [50, 51]. Three different diversity orders can be analyzed: *q* = 0 represents the diversity of all species without overemphasizing their abundances (i.e. an equivalent of species richness), *q* = 1 is the exponential of Shannon’s entropy and represents the diversity of ‘common’ species because it weights each species according to its abundance or frequency. Finally, the order *q* = 2 is the inverse of Simpson’s index and represents the diversity of ‘dominant’ species in the community or, in this study, the assemblage of plant species that were frequently reported as food sources (i.e., rarely exploited plant species are ignored) in the different studies analyzed (Table 1).

To calculate the aforementioned diversity metrics and the sample completeness (i.e., the likelihood that the sample was sufficiently large to detect all the plant species used as food source by each primate species), we used the functions ‘iNEXT’, ‘ChaoRichness’, and ‘estimateD’ of the R package iNEXT [51]. With this same package, we also compared the species richness (*q* = 0) and the diversity of common (*q* = 1) and dominant species (*q* = 2) using rarefaction/extrapolation curves or R/E curves via the function ‘ggiNEXT’. Rarefaction curves are necessary to make appropriate comparisons among communities where the sampling effort is unequal [52, 53], as occurred in this study where the number of study groups was 5, 8, 16, and 34 (Table 2). Using this procedure, it is possible to compute the expected species richness at a standardized sample size and to perform precise extrapolations of the species richness, as the bootstrap samples are added to a pool of previously encountered species [52]. Based on the R/E curves, we used 95% confidence intervals (hereafter 95% C.I.) with the bootstrap procedure to compare the orders of diversity between primate species. Non-overlapping confidence intervals at a particular number of samples indicates statistically significant differences [51, 54]. We estimate the asymptotic α-diversity (i.e., the diversity expected when the R/E curve is asymptotic to the x-axis) using the iNEXT function. We used this procedure to estimate the γ-diversity of plant species in the diet of CR-NHP. For this, we pooled the data for all the study groups and sites.

**Table 2.** Observed and expected diversity of plant species in the diet of the four species of CR-NHP according to the available information on the topic.

| Primate species | Effort<sup>1</sup> | Observed diversity (Hill numbers)<sup>2</sup> |  |  | # gen. | # fam. | #E<sup>3</sup> | Expected diversity (Hill numbers)<sup>4</sup> |  |  |
| --- | --- | --- | --- | --- | --- | --- | --- | --- | --- | --- |
|  |  | 0 | 1 | 2 |  |  |  | 0 | 1 | 2 |
| *Alouatta palliata* | 34 (9000) | 476 (451-501) | 292 | 177 | 247 | 72 | 16 | 854 (748-959) | 431 (400-462) | 203 (186-220) |
| *Ateles geoffroyi* | 8 (5190) | 329 (306-352) | 289 | 245 | 172 | 60 | 4 | 739 (627-850) | 625 (553-696) | 450 (395-505) |
| *Cebus imitator* | 16 (5679) | 236 (221-251) | 191 | 155 | 170 | 67 | 17 | 302 (267-336) | 258 (242-274) | 206 (189-223) |
| *Saimiri oerstedii* | 5 (806) | 183 (161-205) | 170 | 153 | 96 | 48 | 10 | 763 (501-1024) | 645 (465-824) | 408 (322-494) |
| **All** | **61 (20675)** | **868 (829-907)\*** | **520** | **306** | **344** | **87** | **29** | **1740 (1561-1918)** | **774 (726-823)** | **348 (322-374)** |

<sup>1</sup> Sampling effort: number of study groups and total number of sampling hours considering all studies (see Appendix 1 in Chaves *et al.* [49]).
<sup>2</sup> Diversity of species for each Hill' diversity order: 0= species richness, 1= exponential of Shannon entropy or diversity of common species, 2= inverse of Simpson or diversity of dominant species. In the case of species richness (i.e. diversity order 0), the 95% C.I. are showed in parenthesis. \*The observed species richness in diet was 893 species considering the datasets excluded from the analysis.
<sup>2</sup>Number of exotic and/or cultivated plant species used as a food source by each primate species.
<sup>3</sup>Asymptotic diversity estimates according to the non-parametric richness estimators described by [51]. The 95% C.I. for Hill numbers is indicated in parenthesis.

Finally, we compared the Shannon diversity indices for plant species in the diet of each primate species using the Hutchenson t-test [55], using the function ‘multiple_Hutcheson_t_test’ of the R package ecolTest. Because of the large asymmetry in the sampling effort between the four primate species, we used the rarefaction approach to estimate the Shannon indices based on a standardized sample of 5 study groups per primate species.

#### 2.4.2. Plant β-diversity in Diet

We used the dissimilarities between the plant assemblages exploited by primates as indicators of *β* diversity in each of our two habitat types (i.e. TDF and Rainy Forests). As differences between the plant communities can result from two main ecological processes, species turnover and nestedness, we estimated both *β-*diversity components as recommended by Baselga & Orme [56]. For this, we used the function ‘beta-multi’ and ‘beta-sample’ of the R program betapart [57]. In betapart, the species turnover is estimated via the Simpson dissimilarity index (hereafter β_sim_), while the species nestedness is estimated as the nested component of Sørensen dissimilarity (hereafter β_sne_) [56]. Then, we used the function ‘pair-beta’ to perform pairwise comparisons between the plant assemblages exploited by each primate species and the function ‘hclust’ to generate a hierarchical cluster of the pairwise dissimilarities for β_sim_ and β_sne_. To minimize statistical bias due the asymmetric sampling effort, these analyses were based on a rarified random sample of 4 monkey groups in Tropical Dry Forests and 3 groups in Rainy Forests.

#### 2.4.3. Drivers of the Plant Assemblage in the Diet

To determine the influence of the primate species, habitat type, study site, province, forest disturbance level, forest size, forest successional stage, monkey group/population size, study sampling effort (in months), and the interaction between primate species * habitat type on the plant assemblage in diet, we used a permutational multivariable analysis of variance (PERMANOVA), a non-parametric multivariable analysis of variance, as recommended for meta-analyses based in studies with disparate sampling efforts [58]. This method constructs ANOVA-like test statistics from a matrix of resemblances (distances, dissimilarities, similarities) calculated between the sample units, and obtains *p*-values using random permutations of observations among the groups. Furthermore, contrary to traditional parametric multivariable analysis (e.g. MANOVA), PERMANOVA is unaffected by the correlation between variables and is less sensitive to heterogeneity in dispersion [58]. We ran the PERMANOVA via the function ‘adonis2’ of the R package vegan and we set the matrix of Euclidean distances with 999 permutations to estimate the *p*-value. Finally, to determine which levels of the predictor variables differ significantly, we ran post-hoc contrasts using the function ‘pairwise.perm.manova’ of the R package RVAideMemoire. All statistical analyses were performed in R v.4.1.2 [59] and the main R scripts we used are available online (see Appendices 6 and 7 in Chaves *et al.* [49]).

## 3. Results

### 3.1. Alpha and Gamma Plant Species Diversity in the Diet of CR-NHP

Dietary plant species richness varied noticeably within and across records of the four primate species: from 9 to 75 species in *Ap*; 12 to 111 species in *Ag*; 5 to 144 species in *Ci*; and 25 to 92 species in *So* (Table 1). Overall, the total richness of native plant species eaten ranged from 183 species in *So* to 476 species in *Ap* (Table 2, see also Appendix 2 [41]). The four CR-NHP also exploited food items of 29 exotic and/or cultivated plant species, ranging from 4 species in *Ag* to 16 species in *Ap* and *Ci* (Table 2). However, the sampling effort also varied noticeably between species with respect to sampling years and number of study groups. While *Ap* represented 54% of the 61 study groups, *So* represented only 8% (Table 1). Based on our rarefaction curves, the *α*-diversity of plant diet (i.e., diversity order *q*=0) was significantly higher in *Ag* than in the other three primate species, and higher in *So* than in *Ap* and *Ci* (Figure 2; Hutcheson t-test ranged from 4.1 to 26.0, *p* < 0.0001 in all comparisons, see Table S1). The same pattern was also evident in the diversity of common and dominant dietary species (Figure 2).

**Figure 2.**
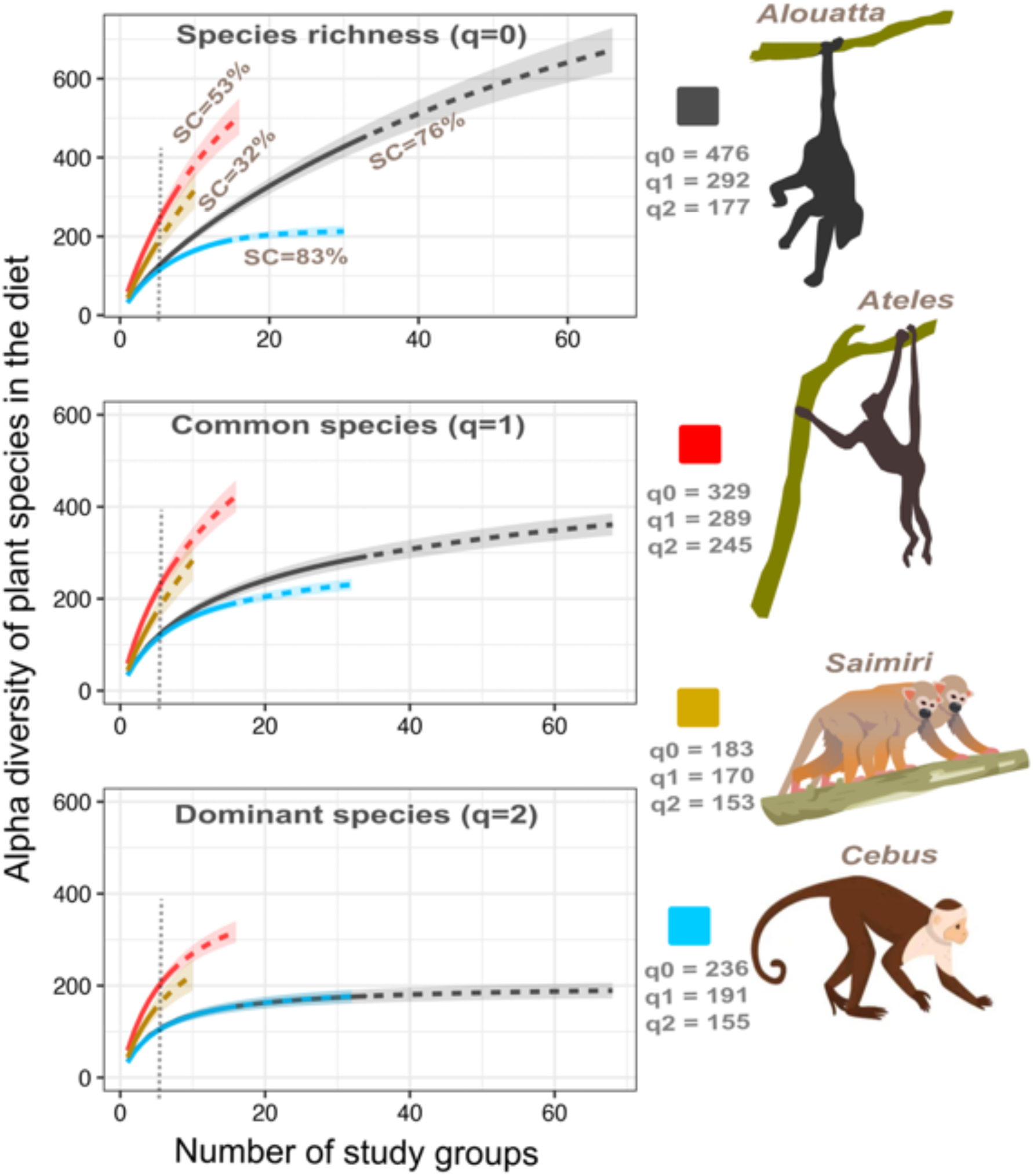
Rarefaction (solid line) and extrapolation (dashed line) curves with 95% confidence intervals comparing the alpha diversity of dietary plants of the four species of CR-NHP for diversity order q=0 (i.e. species richness, top panel), q=1 (i.e., Shannon diversity, middle panel), and q=2 (i.e. Simpson diversity, bottom panel). The reference samples (i.e., the real number of samples analyzed in the package iNEXT) were 34 for *Alouatta*, 8 for *Ateles*, 16 for *Cebus*, and 5 for *Saimiri*. Extrapolation curves are based on the double of the reference sample sizes. Sampling completeness (SC) is indicated for the order q=0. The vertical dotted grey line indicates the rarified sample size (i.e., n = 5 groups for each primate species). The number of plant species exploited according to the diversity order is showed in the legend. Further details on the alpha diversity of plants are provided in Table 2. The 95% C.I. were obtained by bootstrap method based on 200 replications. Non-overlapped 95% C.I. indicate significant differences. See Methods for further details.

Both the standardized plant species richness and the *α*-diversity of common species was similar between *Ap* and *Ci,* but these diversities were higher in *Ap* than in *Ci* when the number of study groups was ≥ 7 (Figure 2). Conversely, the curve for the *α-* diversity of dominant species was flatter for *Ap* and *Ci,* and no difference was detected between these taxa for this diversity metric (Figure 2). The sampling completeness (SC) of the plant species in diet differed between primate species (range = 32%-83%; Kruskal-Wallis test, *H* = 13.6, *p* = 0.003). SC was higher in *Ci* than in *So* (contrast, *p* < 0.05) but no other significant difference was found between the other primate species (contrasts, *p* > 0.05 in all cases, Figure 2). The γ-diversity of plant species in the diet, i.e., pooling the data for the four CR-NHP, was 868 species (including trees, shrubs, palms, epiphytes, parasites, vines, lianas, and terrestrial herbs), distributed across 344 genera and 87 families (Table 2), but the sampling completeness was moderate (i.e. 78%, Figure 3). According to the asymptotic diversity estimates, the expected gamma diversity was 1740 species (range: 1561-1918, Table 2).

**Figure 3.**
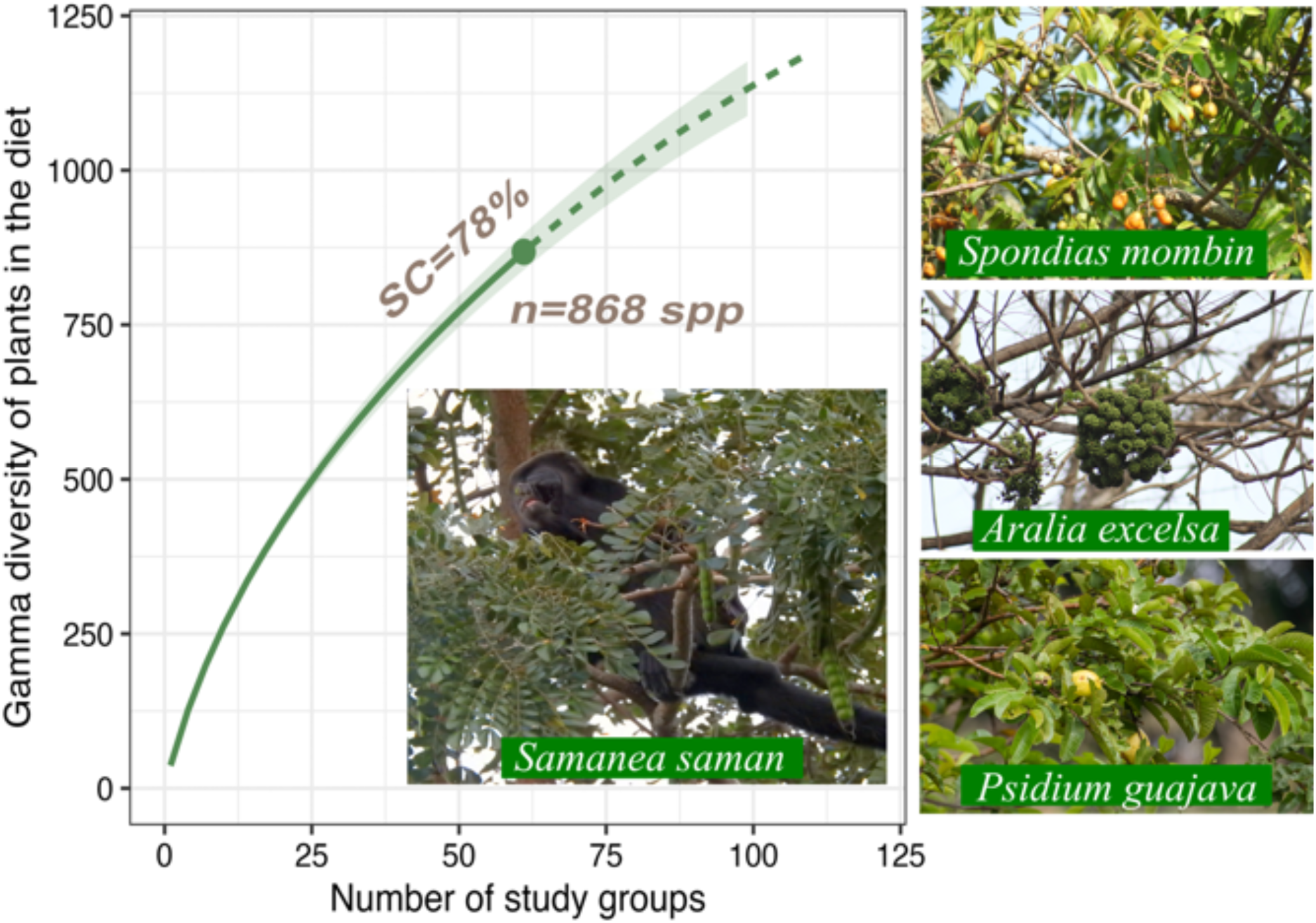
Rarefaction (solid line) and extrapolation (dashed line) curve with 95% confidence intervals for the gamma diversity of plants in diet of the four species of CR-NHP. To perform this diversity profile, the plant species richness (*q* = 0) consumed by the four primate species throughout the study sites were pooled. The extrapolation curve is based on double the reference sample size (i.e. 61 study groups). The 95% confidence intervals were obtained by bootstrapping based on 200 replications. Sampling completeness (SC) and the observed species diversity (n) are indicated in grey bold letters. Images of the single most reported food species for *Alouatta* (i.e. *S. saman*), *Ateles* (i.e. *S. mombin*), *Cebus* (i.e. *A. excelsa*), and *Saimiri* (i.e. *P. guajava*) are shown. Further details on the top ten plant species most frequently reported in the diet of CR-NHP are provided in Table 2.

Finally, in addition to the plant foods, at least ten studies have reported the consumption of animal prey by *Ag, Ci,* and *So.* The consumption of animals was more frequent in *Ci* and *So* which consumed small birds, mammals (including infants of *Cebus imitator*, *Saimiri oerstedii,* and *Sciurus variegatoides* in the case of *Ci*, and *Artibeus watsonii, Uroderma billobatum* and *Vampyresa pusilla* for *So*), amphibians, reptiles, and a large number of invertebrates (i.e. at least 80 taxa, Table S2). Conversely, the consumption of animal prey by *Ap* was extremely rare (i.e. only the caterpillar of *Coenipeta bibitrix*) and was not reported for *Ag* (Table S2).

### 3.2. Most Important Dietary Plant Species

Overall, the three plant species most frequently reported in diet of CR-NHP were, in decreasing order, *Spondias mombin*, *Bursera simaruba*, and *Samanea saman*, while the most important genera were *Ficus*, *Inga*, and *Spondias* (see Appendix 4 in Chaves *et al.* [49]). However, the composition of the top ten reported plant species in the diet of the CR-NHP varied noticeably among species, and animals shared from 0% (*So* vs the other three primate species) to 30% (*Ap* vs *Ag*) of these species (Table 3). Furthermore, the diversity of plant parts eaten varied. *Ap* ingested a diversity of plant parts of the species used (commonly mature and immature fruits and leaves), while *Ag* and *Ci* exploited fewer than two plant parts for most plant species. Squirrel monkeys ingested fruits and nectar from 50% of plant species, but only a single plant part from the other 50% of plant species eaten.

**Table 3.** Top ten plant species most frequently reported in the diet of CR-NHP according to the total number of monkey groups in which each plant species was used as food source.

| Primate species | Plant species<sup>1</sup> | Family | GF<sup>2</sup> | Food items<sup>3</sup> | groups<sup>4</sup> |
| --- | --- | --- | --- | --- | --- |
| *Alouatta palliata*<br>(Ap) | *Samanea saman* | Fabaceae | tree | L, l, lb, F, f, fl | 23 |
|  | *Enterolobium cyclocarpum* | Fabaceae | tree | L, l, F, f, fl | 21 |
|  | *Andira inermis* | Fabaceae | tree | L, l, lb, fl | 19 |
|  | *Spondias mombin* | Anacardiaceae | tree | L, l, lb, F, f, fl | 18 |
|  | *Bursera simaruba* | Burseraceae | tree | L, l, lb, F, f | 17 |
|  | *Astronium graveolens* | Anacardiaceae | tree | L, l, lb, F*, fl, p | 16 |
|  | *Brosimum alicastrum* | Moraceae | tree | L, l, lb, F, f, fl | 16 |
|  | *Maclura tinctoria* | Moraceae | tree | L, l, lb, F, f, fl | 16 |
|  | *Guazuma ulmifolia* | Malvaceae | tree | L, l, F, f, | 14 |
|  | *Handroanthus ochraceus* | Bignoniaceae | tree | L, l, fl | 14 |
| *Ateles geoffroyi*<br>(Ag) | *Spondias mombin* | Anacardiaceae | tree | F, fl | 7 |
|  | *Dilodendron costaricense* | Sapindaceae | tree | F* | 6 |
|  | *Brosimum alicastrum* | Moraceae | tree | L*, F* | 5 |
|  | *B. costaricanum* | Moraceae | tree | F, f | 4 |
|  | *Bursera simaruba* | Burseraceae | tree | l, F* | 4 |
|  | *Cecropia peltata* | Urticaceae | tree | L, l, F*, fl | 4 |
|  | *Garcinia madruno* | Clusiaceae | tree | F | 4 |
|  | *Manilkara chicle* | Sapotaceae | tree | f | 4 |
|  | *Pouteria torta* | Sapotaceae | tree | l, F | 4 |
|  | *Sideroxylon capiri* | Sapotaceae | tree | F, f | 4 |
| *Cebus imitator*<br>(Ci) | *Aralia excelsa* | Araliaceae | tree | L*, F* | 8 |
|  | *Bursera simaruba* | Burseraceae | tree | F* | 8 |
|  | *Muntingia calabura* | Muntingiaceae | tree | F* | 8 |
|  | *Sloanea terniflora* | Elaeocarpaceae | tree | F* | 8 |
|  | *Spondias mombin* | Anacardiaceae | tree | F* | 8 |
|  | *Vachellia collinsii* | Fabaceae | shrub | F* | 8 |
|  | *Genipa americana* | Rubiaceae | tree | F* | 7 |
|  | *Luehea candida* | Malvaceae | tree | F*, fl | 7 |
|  | *Luehea speciosa* | Malvaceae | tree | F*, fl | 7 |
|  | *Ficus* sp. SR | Moraceae | tree | L, l, F* | 7 |
| *Saimiri oerstedii*<br>(So) | *Psidium guajava* | Myrtaceae | tree | F, f | 4 |
|  | *Anacardium excelsum* | Anacardiaceae | tree | F* | 3 |
|  | *Cecropia obtusifolia* | Urticaceae | tree | F* | 3 |
|  | *Conostegia schlimii* | Melastomataceae | tree | F* | 3 |
|  | *Miconia argentea* | Melastomataceae | tree | F* | 3 |
|  | ***Nephelium lappaceum*** | Sapindaceae | tree | F, f | 3 |
|  | *Ochroma pyramidale* | Malvaceae | tree | F*, n | 3 |
|  | *Passiflora vitifolia* | Passifloraceae | vine | F*, n | 3 |
|  | *Symphonia globulifera* | Clusiaceae | tree | F*, n | 3 |
|  | *Vitex cooperi* | Lamiaceae | tree | F* | 3 |
| All primate species | *Spondias mombin* | Anacardiaceae | tree | L, L*, l, F, f, fl, o | 35 |
|  | *Bursera simaruba* | Burseraceae | tree | L, L*, l, F*, fl, o | 29 |
|  | *Samanea saman* | Fabaceae | tree | L, L*, l, F*, F, f, fl, s, o | 27 |
|  | *Brosimum alicastrum* | Moraceae | tree | L, L*, l, F*, F, f, fl, o | 24 |
|  | *Enterolobium cyclocarpum* | Fabaceae | tree | L, L*, l, F*, F, f, fl, s, o | 24 |
|  | *Maclura tinctoria* | Moraceae | tree | L, L*, l, lb, F*, F, f, fl, o | 23 |
|  | *Andira inermis* | Fabaceae | tree | L, L*, l, lb, fl, o | 22 |
|  | *Guazuma ulmifolia* | Malvaceae | tree | L, L*, l, F, F*, f | 21 |
|  | *Muntingia calabura* | Muntingiaceae | tree | L, L*, l, F, F*, fl | 21 |
|  | *Astronium graveolens* | Anacardiaceae | tree | L, L*, l, F*, fl, p, o | 20 |

<sup>1</sup>Plant species are arranged following a decreasing order of primate groups in which each species was exploited and then, by alphabetical order. The entire list of plant species used as food source by each monkey species is provided in the Appendix 4 in Chaves *et al.* [49]. The exotic/cultivated species are enhanced in bold.
<sup>2</sup>Growth form: **tree** = tree > 4 m in height, **shrub** = trees < 4 m in height, **vine** = herbaceous climbers.
<sup>3</sup>Food items consumed by monkeys: **L**= mature leaves, **l** = immature leaves, **lb** = leaf buds, **F** = ripe fruits, **f** = unripe fruits, **fl** = flowers, **s** = seeds, **n** = nectar, **p** = petioles, **o** = others. The asterisk indicates that the maturity stage of the fruits or leaves is unknown.
<sup>4</sup>Number of monkey groups or populations where the consumption of the plant species was recorded.

### 3.3. Plant Species Assemblage Dissimilarity and β-Diversity Components

The number of shared plant species in CR-NHP diets in the TDF ranged from 29 species between *Ap* and *C*i to 44 species between *Ag* and *Ci* , while the number of non-shared species ranged from 33 species between *Ag* and *Ci* to 66 species between *Ap* and *Ci* (Table S3). In Rainy Forests the number of shared plant species in diet ranged from 10 species between *Ci* and *So* to 30 species between *Ag* and *Ap*, while the number of non-shared species ranged from 42 species between *Ag* and *Ci* to 177 species between *Ag* and *So.* In both habitat types species turnover was responsible for most of the dissimilarity in the plant species assemblages exploited by primates (β_sim =_ 0.61 and 0.82, respectively; Figure 4a and c), with the species nestedness explaining only a negligible fraction of the dissimilarity (β_sne =_ 0.03 and 0.05, respectively; Figure 4b and d). Pairwise comparisons of plant β-diversity revealed that in TDF the largest species turnover occurred between *Ap* and *Ci* (β_sim =_ 0.62, Figure 4), while the largest species nestedness was observed between *Ag* and *Ci* (β_sne =_ 0.07). Finally, in Rainy Forests the largest species turnover occurred between *Ap* and *So* (β_sim =_ 0.86), while the largest species nestedness was observed between *Ag* and *Ci* (β_sne =_ 0.17, Figure 4).

**Figure 4.**
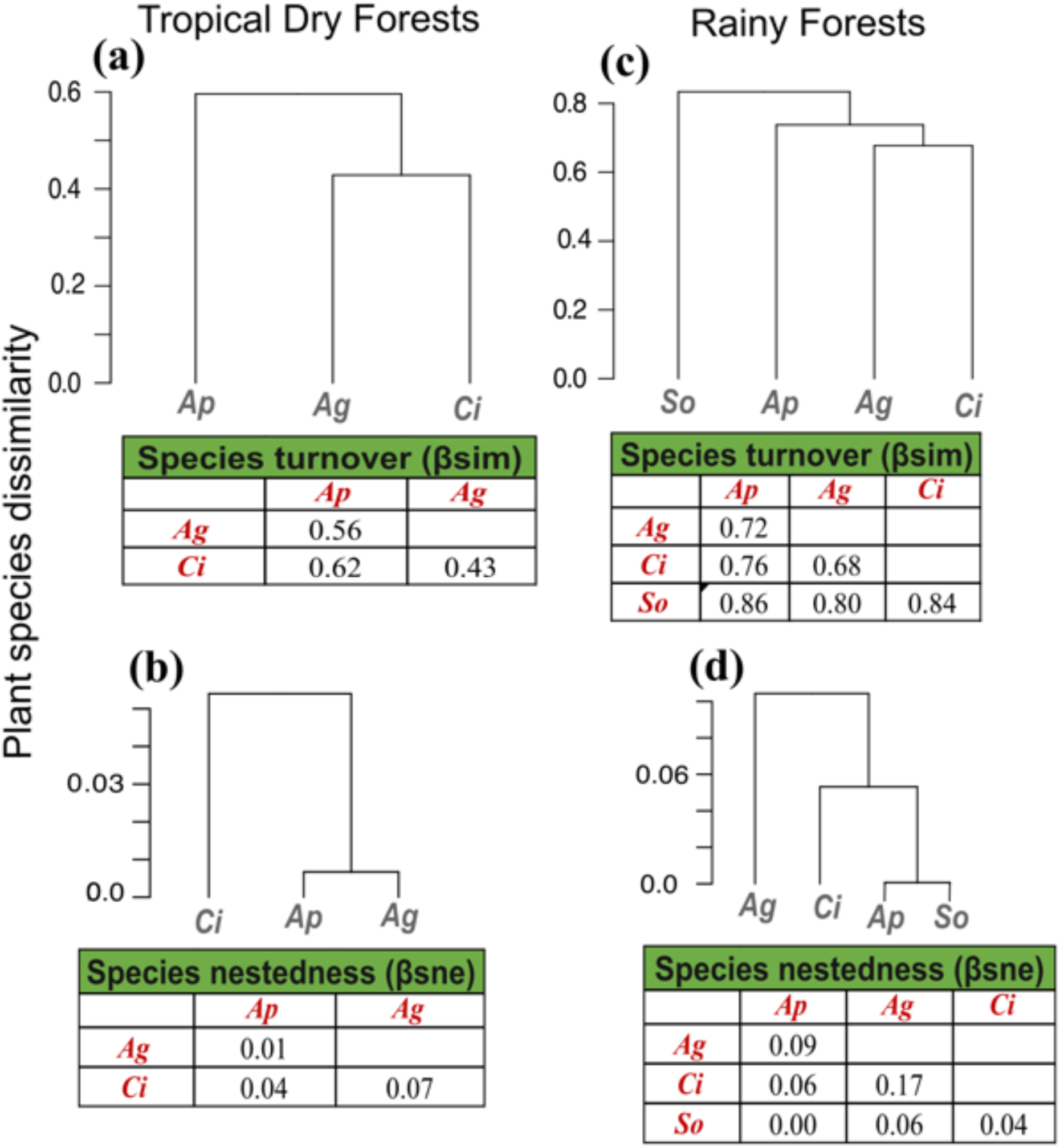
Clustering using average linkage of the β components of plant species dissimilarity between primate species in TDF and Rainy Forests. The component β_sim_ represents the dissimilarity in the plant assemblage due the species replacement or turnover **(a),** while the component β_sne_ represents the plant dissimilarity due the loss/gain of species or nestedness **(b)** in TDF. The same dissimilarity components are showed for Rainy Forests in **(c)** and **(d).** The pairwise comparison values for β_sim_ and β_sne_ are indicted in the tables.

### 3.4. Factors Affecting Plant Assemblages in Diet

The PERMANOVA revealed that three out of five predictor variables significantly influenced plant assemblages exploited by the primates (Table S4). Specifically, we found that the plant composition in the diet was strongly influenced by primate species (F= 1.4, d.f.=3, *p* = 0.002), habitat type (F= 1.8, d.f.=1, *p* = 0.002) and, to a lesser degree, by province (F= 1.4, d.f.=2, *p* = 0.045). However, the rest of variables did not affect the plant assemblages (Table S4). Pairwise comparisons indicated that the plant composition in diet differed significantly between *Ap* and *Ci*, *Ap* and *So*, and *Ci* and *So* (contrasts tests, *p* < 0.05 in all cases), but no other difference was found. Regards the influence of the province, we only detect significant differences between Guanacaste and Limon, and the former province and Puntarenas (contrasts tests, *p* ≤ 0.05 in both cases).

## 4. Discussion

### 4.1. Observed and Expected Diversity of Plants in Diet

Our findings showed that the plant diet of CR-NHP is has considerable species richness. If we consider the expected γ-diversity, the plant assemblage used by these primates was > 1900 plant species, which is consistent with the eclectic feeding behavior observed in other species of Atelidae (e.g. *Alouatta* spp.: [12,13,60], *Ateles* spp. [17, 32]) and Cebidae (e.g. *Cebus* spp. [22,61,62], *Saimiri* spp.: [63, 64], *Sapajus* spp. [11, 65]). Overall, the diverse diets of tropical NHPs are frequently associated with three main factors. Firstly, diet diversification (eating diverse species of fruits, leaves, flowers, invertebrates, etc.) is a frequent behavioral strategy for acquiring essential macronutrients because no single food item typically contains all nutrients [66, 67]. Secondly, diet diversification contributes to diluting the potential negative effects of secondary metabolites present in many plant items (even fruits) that animals ingest, which represent an efficient intoxication-avoiding strategy [68, 69]. Eclectic feeding habits can also be a response to ‘shortage’ of preferred foods which often causes animals to exploit a larger assemblage of fallback foods (e.g. leafy material of climbers growing in forest edges) [70]. Finally, the vegetative constituents of neotropical primate diets in human-modified landscapes includes not only a diverse array of native plants but also dozens of exotic and/or cultivated plants exploited opportunistically (see [9,21,22]), a result we also find in our study. Further studies are necessary to understand if the exploitation of some valuable crops by CR-NHP in the anthropogenic matrix, such as *Persea americana*, *Elaeis guineensis, Syzygium malaccense*, *Spondias purpurea*, and *Mangifera indica* (see Appendix 2 in Chaves *et al*. [49]), result in negative human-NHP interactions, or if crop-raiding behavior is tolerated, as has been reported for *Ap* and *Ag* in Gandoca community, Limon province [71].

In support of our prediction, the α-diversity of plants in diet of *Ag* was higher than in the other three CR-NHP, supporting the hypothesis that the dispersed and unpredictable distribution of fleshy fruits (i.e., the main food item in the diet of *Ag*: [72]) was an evolutionary stimulus to the cognitive adaptations and diet diversification in frugivorous primates [27]. However, the data compiled over 37 years by Melin *et al*. [35], suggests that, at least in some TDFs, the diversity of dietary plants can be higher in *Ci* than in *Ag* (144 vs 89 species) and that the diet of *Ci* can be highly frugivorous, despite *Ci* also consuming a large variety of invertebrates. This study highlights the importance of long-term investigations for improving our understanding of the diet diversity and behavioral flexibility of the four CR-NHP. In addition, our present findings were probably influenced, at least partially, by the fact that all analyzed studies of *Ag* were performed in continuous forests or large well-preserved fragments (the only areas that can support this species), while the studies on the of *Ap* and *So* were performed in continuous and unprotected secondary forest fragments of different sizes (see Table 1). Overall, larger and well-preserved forests are higher-quality habitats for primates because they present a richer food plant assemblage compared with small or medium unprotected secondary forests [4, 73]. It is reasonable to expect that dietary plant diversity reflects these site-specific differences in food availability.

Finally, we believe that the experimental design of some of the studies we analyzed were not robust enough to allow an accurate estimation of the diet diversity (see Table 1), which may influence the α-diversity patterns we obtained. For instance, Chapman [16] reported that in Santa Rosa National Park the diet of *Ap* after a 17-mo study period was only 17 plant species. This is a gross underestimation of the real diet in this habitat, which was later reported to include > 70 plant species [35]. Similarly, studies of *Ci* reported the diet richness was five species [74], and nine species [75]. These studies largely contrast with the plant diet reported for *Ci* in other studies in Santa Rosa National Park (i.e. 144 species [35], 112 species [76]). Our review suggests that these noticeable discrepancies in the plant species richness reported in different studies (even in the same study sites and primate species) are associated with marked differences in the field methods used by each research team, including the observation methods (i.e. instantaneous groups scans, focal animal, *ad libitum* observations, and interviews), the number of observation hours/day/month/year and, specially, the relative knowledge of the monkey observers on the local plant taxonomy. Even when this latter point is crucial to guarantee the data quality and reliability, almost all the reviewed studies omitted the details on how were identified the food plants. If the plant identification was performed by biologists/primatologists with a poor botanical knowledge, the risk of spurious identification and underestimations of plant richness in the diet of CR-NHP increases (particularly in the case of hardly identifiable plants such lianas, vines, and epiphytes).

### 4.2. Most Important Plant Species in Diet

With the caveat that extrapolation of our findings must be considered with caution, due to the lack of systematic feeding records on each food species and the scarcity of diet data (particularly in the case of *Ag* and *So*), our results largely concur with previous studies of the diet of Mesoamerican primates (e.g., [32,60,77,78]. We find that the most important genera in the diets of these animals include large trees such as *Ficus*, *Spondias* and *Brosimum*. This preference is not surprising as these abundant genera produce highly nutritious fleshy fruits, immature leaves, and/or flowers (see [32, 79]). The four CR-NHP shared a low percentage of important plant food species, which is probably explained, at least in part, by the features of the sites they inhabit, in particular the marked contrasts in the structure and composition of plant communities throughout the country [42]. Shifts in the assemblage of top dietary species can also occur due to the existence of particular idiosyncrasies or foraging cultures in some primate groups inhabiting human-modified landscapes [9,21,22]. Our data suggest that this is also the case in CR-NHP. For instance, from the top ten reported plant species in diet of *So*, two can be classified as “cultivated” (Table 3). The fruits of *Nephelium lappaceum* are often exploited and *Psidium guajava* is a highly consumed fruit species frequently grown naturally (or cultivated in some gardens) in open areas throughout the forest edges and the anthropogenic matrix (Ó.M.C. pers. obs.). CR-NHP can be frequently observed foraging in secondary forests and in different elements of the anthropogenic matrix (e.g. subsistence gardens and small agriculture plantations) in Península de Osa, southern Costa Rica [36, 78].

While our findings on the most reported plant species in diet of CR-NHP are preliminary, this plant list provides a useful scientific contribution to the conservation efforts of primates inhabiting severely disturbed and/or urbanized habitats throughout the country. For instance, this type of information has been recently used to elevate management strategies that improve the connectivity (e.g., elaboration of biological corridors, and protection of these tree species in the urban-forest interface) between urban forest remnants inhabiting by *Ap* in different highly touristy and urbanized areas of Guanacaste, Costa Rica [40,41,47, 80]. In this respect, the most promising food species for improving the forest connectivity through reforestation are likely rapid-growing and/or pioneer native species such as *Ficus* spp., *Spondias mombin*, *Bursera simaruba*, and *Muntingia calabura* (see Appendix 2 and 4 in [49] for further information on the plant food species). Reforestation including these species, together with the installation of artificial bridges and environmental education, can be powerful allies to prevent or mitigate the all too frequent events of electrocution and vehicle-collisions of primates in Guanacaste and other regions of Costa Rica [41,47,80].

### 4.3. Plant β-diversity and Factors affecting the Plant Composition in Diet

As predicted, our findings strongly suggest that plant species turnover was the main mechanism responsible for the dissimilarities between the dietary plant assemblages of CR-NHP in both Tropical Dry Forests and Rainy Forests. Interestingly, we found that plant species replacement increased in the latter habitats. This pattern is likely explained, at least in part, by the fact that the Rainy Forests not only included four different HLZ, but that the study sites also varied in size, successional stage, protection status, and probably, forest cover. Therefore, even though most of the reviewed papers did not provided data on the site-specific plant composition, the high dissimilarity in the diets of CR-NHP may reflect the plant species replacement between sites and/or the contrasting feeding and digestive strategies of each primate species (e.g., Milton [27, 30]). Our findings also showed that the habitat type and the province were the best predictors of plant assemblage in the diet of these animals. Overall, in many tropical forests, the β-diversity of tree species is mainly associated with species turnover because of spatial variability in climatic and biotic conditions [5, 81]. This environmental variability actively promotes plant species replacement from one habitat type and/or province to another due to processes such as dispersal limitation and speciation [26, 81] and diverse anthropogenic activities that alter the plant structure and quality of primate habitats (e.g. extensive agricultural practices, urbanization, and cattle ranching: [19, 20]).

Even though the forest cover of Costa Rica is relatively small compared with most South American countries, differences in the plant community composition tend to increase with distance. Changes in plant composition are notable from the northern Tropical Dry Forests to the southern Tropical Rain Forests [42] and even within a single region [82]. Therefore, due to the inter-site variation in plant composition, CR-NHP adjust their diets to the available plant foods [2,70,83], which result in large plant species turnover in the diet. Finally, the contrasting foraging and digestive strategies observed between primate species (e.g., *Ap* vs *Ag*: [27, 30]) may also influence the plant composition in diet of each CR-NHP.

## 5. Conclusions

Despite the scarcity of information on the topic (particularly for *So* and *Ag*) this meta-analysis supports the conclusion that CR-NHP have highly diverse and eclectic diets. Taken together, these primates used at least 868 plant species (including 29 exotic and/or cultivated species). Rarefaction curves we used in our analysis suggest higher α-diversity of plants in the diet of *Ag* compared with the other three primate species, supporting previous studies on the importance of unpredictable fleshy fruits as a selective pressure for diet diversification [27]. Similarly, we also find support that plant species turnover was the main ecological mechanism explaining the dissimilarity of plant assemblages in diet of the CR-NHP. This was largely expected, considering that vegetation assemblages (i.e., the food source available for primates) can vary drastically throughout the country [42, 46]. The data we have compiled improves our understanding on the feeding ecology and the dietary flexibility of CR-NHP and provides valuable scientific input regarding the design and implementation of management strategies.

At the same time, we recognize that most of the patterns in dietary diversity we reported here could change noticeably if we took into consideration the entire diet (i.e. considering both plant and animal items consumed by primates). Unfortunately, this type of analysis is not possible because, to date, most records of animal items exploited by CR-NHP are anecdotal, incomplete, and strongly biased to *Ci* and *So* (Table S1). Recent studies in Para state, Brazil, indicate that even in two highly frugivorous *Ateles* species, the consumption of animal material occurs frequently because spider monkeys intentionally select larvae-infected fruits as possible strategy to satisfy the protein demand [84], and the same situation may be true in the CR-NHP (and in most of the 179 recognized Neotropical primates: [39]). Therefore, despite the multiple challenges associated with the study of animal consumption by primates (e.g., difficulty to determine the larval content of ripe fruits ingested, invertebrate complex taxonomy, and rarity of some hunting events of small and medium vertebrates), further studies on this topic are crucial to understand the diet diversity of Neotropical primates. In this sense, we suggest the use of advanced molecular techniques, such as metagenomic analysis of fecal samples, an approach that has been used with success in different Neotropical primates [62, 85]. These techniques can also complement the observational analysis of plant consumption, particularly when plant species cannot be identified by the observers due to challenges in the field.

### Supplementary Materials

The following supporting information can be downloaded at: https://www.mdpi.com/article/XXXX. Table S1: Results of the Shannon-Wiener indexes comparison for the plant dietary diversity for Costa Rican non-human primates using a rarified sample; Table S2: Animal taxa reported in the diet of CR-NHP; Table S3: Shared and non-shared plant species in the diet of the four Costa Rican non-human primates; Table S4: Results of the PERMANOVA.

### Author Contributions

Conceptualization, formal analysis, writing-original draft preparation, visualization, supervision, project administration, Ó.M.C.; data collection via literature review and/or fieldwork, Ó.M.C., J.C.Q, V.C.M., I.A.R., D.S., P.R.H., C.C.; data curation, E.C.M., Ó.M.C., V.C.M., I.A.R.; writing—review and editing, Ó.M.C., V.C.M., I.A.R., P.R.H., E.C.M., D.S., C.C., A.D.M; funding-acquisition, Ó.M.C. All authors have read and agreed to the published version of the manuscript.

### Funding

This research was funded by the Vicerrectoría de Investigación of the Universidad de Costa Rica, project # 908-C1611, the Programa de Voluntariado-UCR.

### Institutional Review Board Statement

Not applicable.

### Informed Consent Statement

Not applicable.

### Data Availability Statement

All the datasets associated with this study are available at: https://doi.org/10.6084/m9.figshare.21785588.v1. Appendix 1: Detailed list of the studies on the diet of CR-NHP reviewed in this meta-analysis; Appendix 2: Database on the food species exploited by the four primate species; Appendix 3: PDFs of the papers included int Table 1, Appendix 4: List of the main plant species, orders and families used by monkeys as food source; Appendix 5: R scripts on the analysis and main results of the diversity analysis in iNEXT; Appendix 6: R scripts and results of the PERMANOVA assessing the influence of the predictor variables on the food plant assemblage. These datasets are also cited in the text [41].

## Acknowledgments

We thank Elder Gómez, Roy J. Vallejos, Adonis Gamboa, Kenneth Barrantes, and Josue Umaña for field and lab assistance. To Owen H. Cerdas (Tamarindo Park) and Jose J. Somarribas (Finca Josema) for giving us permission to conduct this research in their properties in Tamarindo and 27 de Abril. Marco Retana provided valuable logistical support to locate the howler monkey groups in Tamarindo and 27 de Abril. The Programa de Voluntariado-UCR, led by Lupita Abarca Espeleta, provided financial support to the maintenance of the volunteers during 2021 and 2022. The botanist Mario Blanco collaborated in the identification of the plant species used by howler monkeys in Santa Cruz and in the elaboration of the reference collection of plants at the Herbario Luis Fournier. We thank the administration team in Santa Rosa National Park, Sector Santa Rosa, for supporting our research and assistance with permits and logistics over many decades, especially Roger Blanco and Maria Martha Chavarria. We are grateful to the many dedicated students and assistants that have worked on this project with special thanks to Saúl Cheves Hernandez, Adrian Guadamuz Chavarria, Evin Murillo Chacon, and Ronald Lopez Navarro. Finally, we thank to two anonymous reviewers for their useful commentaries and recommendations to improve the early version of this manuscript.

## Conflicts of Interest

The authors declare no conflict of interest.

## Notes

https://doi.org/10.6084/m9.figshare.21785588.v1

### Competing Interest Statement

The authors have declared no competing interest.

### Summary of Updates

In human-modified tropical landscapes, the survival of arboreal vertebrates, particularly primates, depends on their plant dietary diversity. Here, we assessed diversity of plants included in the diet of Costa Rican non-human primates, CR-NHP (i.e. Alouatta palliata palliata, Ateles geoffroyi, Cebus imitator, and Saimiri oerstedii) inhabiting different habitat types across the country. Specifically, we assessed by analyzing 37 published and unpublished datasets: (i) richness and dietary Alpha-plant diversity, (ii) the β-diversity of dietary plant species and the relative importance of plant species turnover and nestedness contributing to these patterns, and (iii) the main ecological drivers of the observed patterns in dietary plant . Diet data were available for 34 Alouatta, 16 Cebus, 8 Ateles, and 5 Saimiri groups. Overall dietary plant species richness was higher in Alouatta (476 spp.), followed by Ateles (329 spp.), Cebus (236 spp.), and Saimiri (183 spp.). However, rarefaction curves showed that Alfa-diversity of plant species was higher in Ateles than in the other three primate species. The γ-diversity of plants was 868 species (95% C.I.=829-907 species). The three most frequently reported food species for all CR-NHP were Spondias mombin, Bursera simaruba, and Samanea saman. In general, plant species turnover, rather than nestedness, explained the dissimilarity in plant diet diversity (βsim> 0.60) of CR-NHP. Finally, primate species, habitat type (life zone and disturbance level) and, to a lesser degree, sampling effort were the best predictors of the dietary plant assemblages. Our findings suggest that CR-NHP diets were diverse, even in severely-disturbed habitats.

