## Supplementary Material_f for "Plant diversity in the diet of Costa Rican primates in contrasting habitats: a meta-analysis"

This is the supplementary material associated to the following scientific article:

**Supplemental Data:**

**Collection Methods Used in Unpublished Studies**

Chaves *et al*. (unpl. data) collected data on the diet of three habituated *Alouatta palliata* groups during a 16-month period (Jun 2021-Sept 2022), that included the study period of this meta-analysis (Table 1). In this investigation, ÓMC, VMC, and JC followed the groups 1-2/days per month from dawn to dusk. The behavioral records were collected with the aid of high-definition binoculars (Nikon® Prostaff 5, 10 x 52) via instantaneous scans every 15 min during a 5 min period and, during the inter-scan period, 5-min focal animal samples were collected when some individual was feeding. Riba-Hernández & Stoner (unp. data) and Stoner & Riba-Hernández (unpl. data) used the same field methods described in Stoner *et al*. (2005), i.e. 2-min focal animal observations, to study the foraging behavior of three free-ranging groups of *A. palliata*. Finally, in the forest fragment of the Danta Corcovado Lodge, Guadalupe, Península de Osa (N8.62103, W83.47594), Solano-Rojas (unpl. data) collected data on the diet of a habituated free-ranging group of *Saimiri o. oerstedii* during a 8-mo period (Dec 2018-July 2019) distributed in 62 sampling days.

The fieldwork in the investigation of Chaves *et al*. (unpublished. data) was performed with the approval of the Comité Institucional de Cuidado y Uso de Animales of the Universidad de Costa Rica (Ethic authorization #CICUA-071-2021), the Comisión Institucional de

Biodiversidad (authorization #CBio-4-2021), and the Sistema Nacional de Áreas de Conservación, SINAC (permission # ACT-OR-DR-110-2021).

### Comparison of alpha-diversity of species in diet using the Shannon-Wiener index

Using a standardized number of study groups per primate species (i.e. 5 study groups), the plant species richness in diet was 217 species for *Ateles geoffroyi* (Shannon index, $H$ = 2.30), 183 for *Saimiri oerstedii* ($H$ = 2.23), 145 for *Cebus imitator* ($H$ = 2.11), and 66 species in *Alouatta palliata* ($H$ = 1.80). Consequently, the Shannon indices differed significantly among primate species (see Table S1).

**Table S1.** Results of the Shannon-Wiener indexes comparison for the plant dietary diversity for Costa Rican non-human primates using a rarified random number of monkey groups (i.e. 5 groups/primate species).

| Comparison[1] | Shannon-Wiener index[1] | Hutcheson t-test | d.f. | P-value |
|---|---|---|---|---|
| ***Ateles*** *vs Alouatta* | 2.30 vs 1.80 | 25.9 | 169 | <0.0001 |
| ***Ateles*** *vs Cebus* | 2.30 vs 2.11 | 10.0 | 418 | <0.0001 |
| ***Ateles*** *vs Saimiri* | 2.30 vs 2.23 | 4.1 | 472 | <0.0001 |
| ***Saimiri*** *vs Alouatta* | 2.23 vs 1.80 | 21.7 | 181 | <0.0001 |
| ***Saimiri*** *vs Cebus* | 2.23 vs 2.11 | 6.1 | 412 | <0.0001 |
| ***Cebus*** *vs Alouatta* | 2.11 vs 1.80 | 14.6 | 215 | <0.0001 |

[1]The species with higher plant diversity in diet is enhanced in bold.

[2]The Shannon diversity index for each primate species is indicated

**Table S2**. Animal taxa reported in the diet of Costa Rican primates up to October 2022.

| Species/Mspp. | *A. palliata* | *C. imitator* | *S. oerstedii* | Reference |
|---|---|---|---|---|
| **VERTEBRATES** | | | | |
| **Mammalia** | | | | |
| Primates | | | | |
| *Cebus imitator* | | X | | Nishikawa *et al*. (2020) |
| *Saimiri oerstedii* | | X | | Chaves (unp. data) |
| Didelphidae | | | | |
| *Caluromys derbianus* (tail) | | X | | Chaves (unp. data) |
| Procyonidae | | | | |
| *Nasua narica* | | X | | Fedigan (1990) |
| Rodentia | | | | |
| *Sciurus variegatoides* | | X | | Chaves (unp. data) Fedigan 1990 |
| Chiroptera | | | | |
| *Artibeus watsoni* | | | X | Boinski & Tim (1985) |
| bat Msp1 | | X | X | Boinski (1986), Fedigan (1990) |
| **Aves** | | | | |
| bird Msp1 | | X | X | Boinski (1986), Fedigan (1990) |
| various birds | | X | | Fedigan (1990) |
| bird eggs | | | | Fedigan (1990) |
| **Reptilia** | | | | |
| snails | | X | X | Boinski (1986), Fedigan (1990) |
| lizards | | X | X | Boinski (1986), Fedigan (1990) |
| *Anolis biporcatus* | | X | | Chaves (unp. data) |
| *Ctenosaura similis* (tail) | | X | | Chaves (unp. data) |
| *Iguana iguana* (tail) | | X | | Chaves (unp. data) |
| **Amphibia** | | | | |
| frogs Mspp | | X | X | Boinski (1986), Melin (unp. data) |
| **INVERTEBRATES** | | | | |
| **Arachnida** | | | | |
| spider Msp1 | | X | X | Boinski (1986), Melin (unp. data) |
| **Abranchiata** | | | | |
| Lumbricidae | | | X | Boinski (1986) |
| **Pulmonata** | | | | |
| flatworms | | | X | Boinski (1986) |
| earthworms | | | X | Boinski (1986) |
| **Insecta** | | | | |
| insect eggs | | | X | Boinski (1986), Melin (unp. data) |

| | | | | |
|---|---|---|---|---|
| Formicidae | | | | |
| *Pseudomyrmex* sp. | | X | | McCabe (2005) |
| *Pseudomyrmex flavicornis* | | X | | Freeze (1976) |
| Hemiptera | | X | X | Boinski (1986), Melin *et al.* (2010) |
| Homoptera | | | X | Boinski (1986) |
| Hymenoptera | | | | |
| ants, Boinski | | | X | Boinski (1986), Melin et al 2010 |
| wasp, Boinski | | | X | Boinski (1986), Melin e*t al.* (2010) |
| Blattodea | | | | |
| cockroach Msp1 | | X | X | Boinski (1986), Melin *et al.* (2010) |
| insect Msp12 | | X | | Boinski (1986) |
| insect Msp13 | | X | | Boinski (1986) |
| Coleoptera | | | | |
| beetles | | X | X | Boinski (1986), Melin *et al.* (2010) |
| Diptera | | | | |
| flies | | | X | Boinski (1986) |
| Lepidotera | | | | |
| Lepidoptera Msp1 | | X | X | McCabe (2005) |
| Erebidae | | | | |
| *Coenipeta bibitrix* (caterpillar) | X | | | Azofeifa (2022) |
| Orthoptera | | | | |
| Orthoptera Msp1 | | X | X | Boinski (1986), Melin *et al.* (2010) |
| Tettigoniidae | | | | |
| katydid Msp1 | | X | X | Boinski (1986), Melin *et al.* (2010) |
| Termitidae | | | | |
| Termitidae | | X | X | Boinski (1986), Melin *et al.* (2010) |
| **Vespidae** | | | | |
| *Polybia* sp | | | X | Boinski & Tim (1985) |
| Insects no identified | | | | |
| insects (8 Mspp.) | | X | | Melin *et al.* (2014) |
| insects (48 Mspp.) | | X | | Mosdossy *et al.* (2015) |
| **Total = 13 species + 81 Mspp.** | **1** | **71** | **23** | |

**Table S3.** Shared and non-shared plant species in the diet of the four Costa Rican non-human primates in Tropical Dry Forests and Rainy Forests. For this analysis we only considered the studies ≥ 6 months in duration. *Ap* = *Alouatta palliata palliata*, *Ag* = *Ateles geoffroyi*, *Ci*= *Cebus imitator*, *So* = *Saimiri oerstedii.* The data are based in a standardized sample of 4 monkey groups in Tropical Dry Forests and 3 groups in Rainy Forests. **S. oerstedii* do not occurs in Tropical Dry Forest.

| Tropical Dry Forests* | | | | Rainy Forests | | | | |
|---|---|---|---|---|---|---|---|---|
| Shared plant species in diet | | | | Shared plant species in diet | | | | |
| | *Ap* | *Ag* | *Ci* | | *Ap* | *Ag* | *Ci* | *So* |
| *Ap* | 95 | 41 | 29 | *Ap* | 106 | 30 | 15 | 15 |
| *Ag* | 41 | 98 | 44 | *Ag* | 30 | 198 | 20 | 21 |
| *Ci* | 29 | 44 | 77 | *Ci* | 15 | 20 | 62 | 10 |
| | | | | *So* | 15 | 21 | 10 | 107 |
| Non-shared plant species in diet | | | | Non-shared plant species in diet | | | | |
| | *Ap* | *Ag* | *Ci* | | *Ap* | *Ag* | *Ci* | *So* |
| *Ap* | 0 | 57 | 48 | *Ap* | 0 | 168 | 47 | 92 |
| *Ag* | 54 | 0 | 33 | *Ag* | 76 | 0 | 42 | 86 |
| *Ci* | 66 | 54 | 0 | *Ci* | 91 | 178 | 0 | 97 |
| | | | | *So* | 91 | 177 | 52 | 0 |

**Table S4.** Results of the PERMANOVA assessing the influence seven predictors on the plant species assemblage in diet of Costa Rican primates. Significant effects are enhanced in bold.

| Predictor variable | d.f. | Sum of squares | *F* | $R^2$ | *p* |
|---|---|---|---|---|---|
| Primate species | 3 | 8.2 | 1.4 | 0.15 | **0.002** |
| Habitat type (i.e. TDF and Rainy Forest) | 1 | 3.5 | 1.8 | 0.06 | **0.002** |
| Province | 2 | 5.3 | 1.4 | 0.09 | **0.045** |
| Group size | 1 | 2.6 | 1.4 | 0.05 | 0.052 |
| Study site | 10 | 23.5 | 1.2 | 0.42 | 0.075 |
| Primate species*HLZ | 1 | 2.3 | 1.2 | 0.04 | 0.14 |
| Sampling effort | 1 | 2.2 | 1.1 | 0.04 | 0.24 |
